# Identifying age cohorts responsible for *Peste des petits ruminants* virus transmission among sheep, goats, and cattle in northern Tanzania

**DOI:** 10.1101/2019.12.19.882233

**Authors:** C. M. Herzog, W. A. de Glanville, B. J. Willett, I. M. Cattadori, V. Kapur, P. J. Hudson, J. Buza, E. S. Swai, S. Cleaveland, O. N. Bjørnstad

**Affiliations:** Center for Infectious Disease Dynamics, Pennsylvania State University, State College, Pennsylvania, USA; Institute of Biodiversity, Animal Health and Comparative Medicine, University of Glasgow, Glasgow, UK; MRC-University of Glasgow Centre for Virus Research, Glasgow, UK; Nelson Mandela African Institute of Science and Technology, Arusha, Tanzania; Department of Veterinary Services, Ministry of Livestock and Fisheries, Dodoma, United Republic of Tanzania

## Abstract

*Peste des petits ruminants* virus (PPRV) causes a contagious disease of high morbidity and mortality in global sheep and goat populations and leads to approximately $2 billion USD in global annual losses. PPRV is currently targeted by the Food and Agricultural Organization and World Animal Health Organization for global eradication by 2030. To better control this disease and inform eradication strategies, an improved understanding of how PPRV risk varies by age is needed. Our study used a piece-wise catalytic model to estimate the age-specific force of infection (FOI, per capita infection rate of susceptible hosts) among sheep, goats, and cattle from a cross-sectional serosurvey dataset collected in 2016 in Tanzania. Apparent seroprevalence rose with age, as would be expected if PPRV is a fully-immunizing infection, reaching 53.6%, 46.8%, and 11.6% (true seroprevalence: 52.7%, 52.8%, 39.2%) for sheep, goats, and cattle, respectively. Seroprevalence was significantly higher among pastoral animals than agropastoral animals across all ages, with pastoral sheep and goat seroprevalence approaching 70% and 80%, respectively, suggesting endemicity in pastoral settings. The best fitting piece-wise catalytic models included merged age groups: two age groups for sheep, three age groups for goats, and four age groups for cattle. However, the signal of these age heterogeneities was weak, with overlapping confidence intervals around force of infection estimates from most models with the exception of a significant FOI peak among 2.5-3.5 year old pastoral cattle. Pastoral animals had a higher force of infection overall, and across a wider range of ages than agropastoral animals. The subtle age-specific force of infection heterogeneities identified in this study among sheep, goats, and cattle suggest that targeting control efforts by age may not be as effective as targeting by other risk factors, such as management system type. Further research should investigate how specific husbandry practices affect PPRV transmission.

**Author Summary:** Age differences in transmission are important for many infections, and can help target control programs. We used an age-structured serosurvey of Tanzanian sheep, goats, and cattle to explore *peste des petits ruminants* virus transmission. We estimated rate at which susceptibles acquire infection (force of infection) to determine which age group(s) had the highest transmission rates. We hypothesized that an age-varying model with multiple age groups would better fit the data than an age constant model and that the highest transmission rates would appear in the youngest age groups. Furthermore, we hypothesized evidence of immunity would increase with age. The data supported our hypothesis at the species level and the best fitting models merged age groups: two, three, and four age group models were best for sheep, goats, and cattle, respectively. The highest rates occurred among younger age groups and evidence of immunity rose with age for all species. In most models, confidence interval estimates overlapped, but there was a significant FOI peak among 2.3-3.5 year old pastoral cattle. Importantly, these data indicate that there is not sufficient evidence to support targeted control by age group, and that targeted control based on production system should be more effective.

## Introduction

*Peste des petits ruminants* virus (PPRV), or small ruminant morbillivirus (SRMV) is a socio-economically important, highly infectious virus that causes high morbidity and mortality among sheep and goat populations worldwide. PPRV is present in Africa, the Middle East, Asia, and Turkey and currently impacts 80% of the world’s sheep and goat population [1], causing an estimated $1.45-2.1 billion USD in global annual losses due to mortality, impaired production, and treatment for infected animals [1]. Livestock keepers in these regions rely heavily on sheep and goats for their livelihoods, as they are a source of meat, milk, and income. Household herd losses due to PPRV contribute to global poverty and food insecurity. In 2015, the Food and Agricultural Organization (FAO) and World Animal Health Organization (OIE) launched a global campaign to eradicate PPRV by 2030. Although the available, affordable vaccine demonstrates protection for up to 3 years [2], the large, global small ruminant population turns over rapidly, increasing the difficulty of control strategies such as blanket mass vaccination. To better target eradication control efforts, FAO also launched a global PPRV research network in 2018 [3] with the goal of aligning research efforts to inform strategies for PPRV eradication. One such research effort highlighted was a need to determine how PPRV transmission patterns vary by age, including identifying the appropriate age cohorts at which PPRV vaccination can be performed efficaciously and the age at which maternal immunity falls beneath a protective threshold (and what the value of the protective threshold is) [4].

The role of age in vulnerability to disease transmission is an important consideration that often leads public health and animal health experts to focus their control efforts on specific subsets of populations [5]. For many well-known infectious diseases, transmission risk varies by age. For example, measles, mumps, and rubella, became known as childhood diseases in the pre-vaccine era as the force of infection (FOI, per capita rate of infection of susceptible hosts) was greatest during early childhood [6,7] due to age-stratified mixing and an adult population protected from previous exposure. Sexually transmitted infections typically have higher incidence and FOI after the age of sexual maturity among more sexually active individuals. Other infections, including influenza or bacterial infections, may impact any age group but have a greater disease impact on those who are immunocompromised – often the youngest and oldest members of a population [8]. In animals, evidence of age-dependent patterns in susceptibility and transmission have been observed for feline leukemia virus in cats and bovine tuberculosis in bison [9–11]. In these infections, young animals were more susceptible to infection and seropositivity increased in the population with age. These age-related infection patterns are influenced by mixing patterns, which are likely to be different among wild animals, domesticated animals in different production systems, and children, who experience age-stratified mixing at school. Nevertheless, given that age-related patterns are seen among both human and animal diseases, calculating the FOI across age cohorts is a useful approach that can reveal which age group is responsible for most of the onward transmission. Prevention and control efforts can then be targeted to the appropriate age cohorts, reducing transmission and lowering control costs when compared with mass interventions, assuming the cost of identifying target groups is not prohibitive [12].

PPRV research to date has demonstrated that young animals may be protected for the first 3-5 months of life by maternal immunity [13–17], but once protection wanes young animals appear to be the most severely affected age group in terms of susceptibility and mortality [16–18]. The majority of published studies (compiled in S1Text) report age as a significant risk factor for PPRV seroconversion and that PPRV seroprevalence increases with animal age; however, there are also several studies that concluded age was not a significant risk factor so further clarification is needed regarding the impact of age on PPRV transmission.

To understand the role of age in PPRV transmission and to inform strategies for PPR eradication by vaccination, this study aims to identify which age cohorts are responsible for PPRV transmission using a cross-sectional serosurvey data of sheep, goats, and cattle managed together in the same households under differing production systems across 20 villages in northern Tanzania. Earlier work in this population [19] did not assess age variation in transmission. We predict that seroprevalence will increase by age group for each species, as would be expected if PPRV is an endemic, fully-immunizing (i.e. providing life-long protection) infection. We use a piece-wise catalytic model framework to calculate age-varying FOI estimates. Our *a priori* set of hypotheses for the piece-wise catalytic model are represented schematically in Fig 1. We postulate that an age-varying model (Figs 1B-1D) will be a better fit to serosurvey data than an age constant model (Fig 1A). Furthermore, we predict that the highest FOI estimates will appear in the youngest age groups, shortly after waning of maternal immunity, consistent with a hypothesis of horizontal transmission by socialization at age of weaning and independence around a few months of age, or, in the case of sheep and goats, reproduction starting shortly before one year of age (Fig 1B). It will be possible to detect, but not separate, these two events in sheep and goats, as animals in the data set are all six months or older and the dentition-based age groups do not provide fine enough age resolution. As a precaution, models without the first age group were investigated in case of extended maternal immunity beyond published estimates. The hypothesized transmission patterns for these models are presented in Figs 1C-1D, and include an abbreviated FOI peak in the youngest age groups.

**Figure 1.**
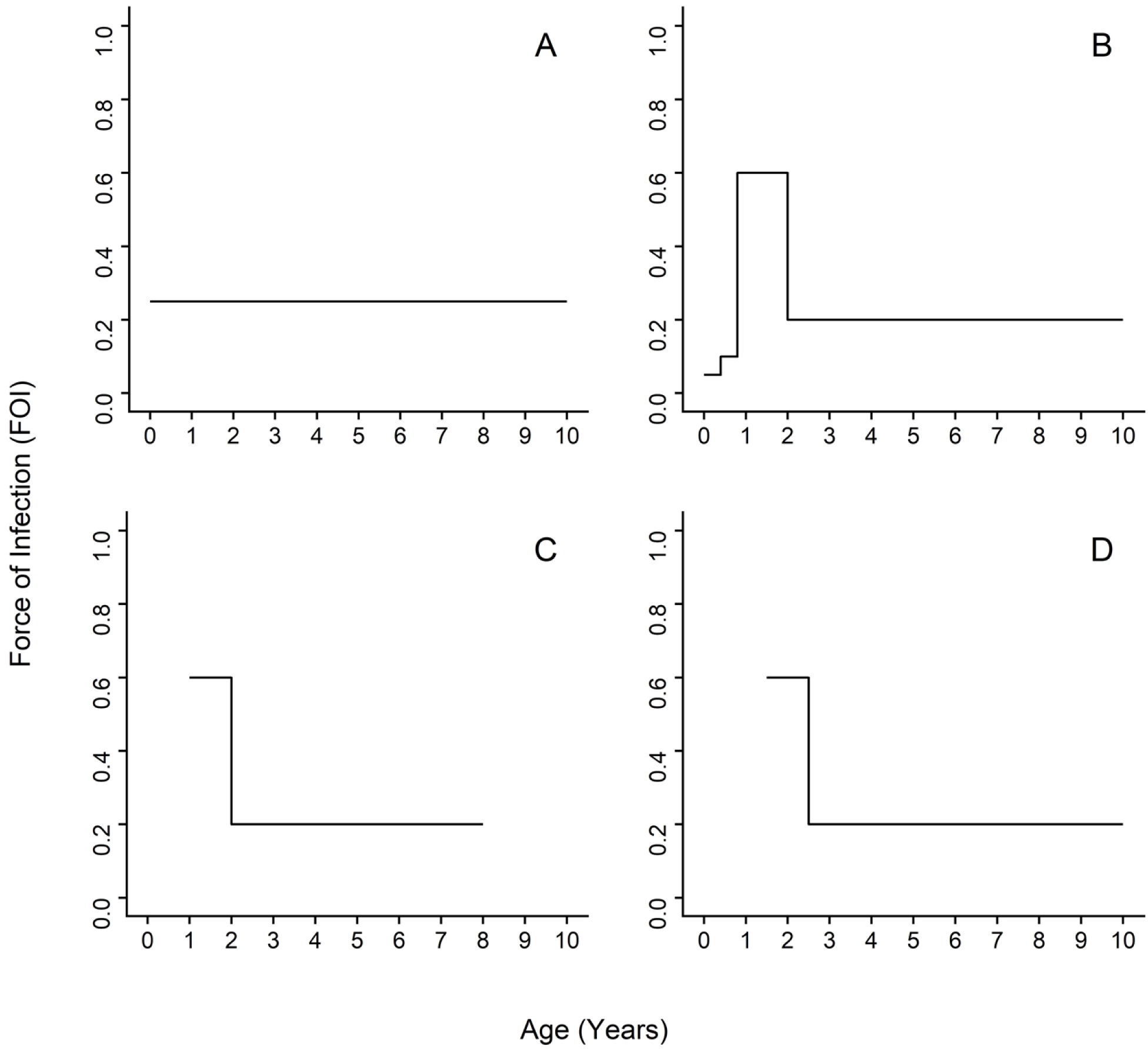
Schematic representation of possible age-specific force of infection PPRV transmission patterns. Possible transmission routes include A) constant from birth due to environmental exposure, B) horizontal transmission during routine social activities at age of independence and then at time of breeding, C) expected pattern among sheep and goats in five age group data set dominated by horizontal transmission due to breeding, D) expected pattern among cattle in five age group data set dominated by horizontal transmission due to socialization at age of independence.

## Results

Demographic characteristics of the 7,496 samples analyzed were published previously [19], a subset of which are included in Table S1. Overall, there were 1,869 (24.9%), 757 (10.1%), 666 (8.9%), 483 (6.4%), 3,307 (44.1%), and 414 (5.5%) animals in the age groups 1-6, respectively (Fig 2, Table S2). For each species, the largest number of sampled animals was found in the fifth age group (full mouth with no wear), followed by the first age group (temporary teeth).

**Figure 2.**
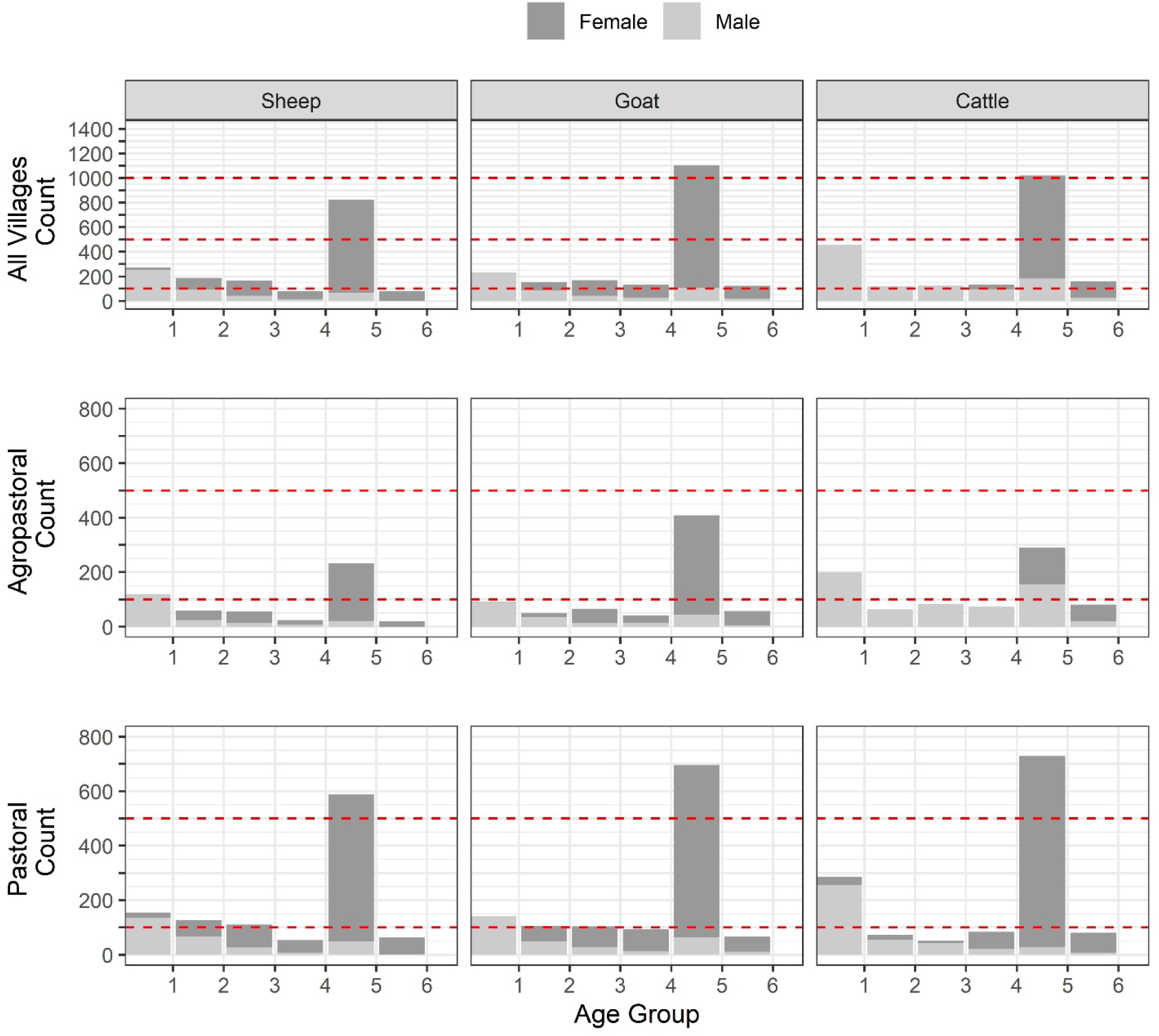
Dentition-based age group distribution by species, sex, management system.

**Figure 3.**
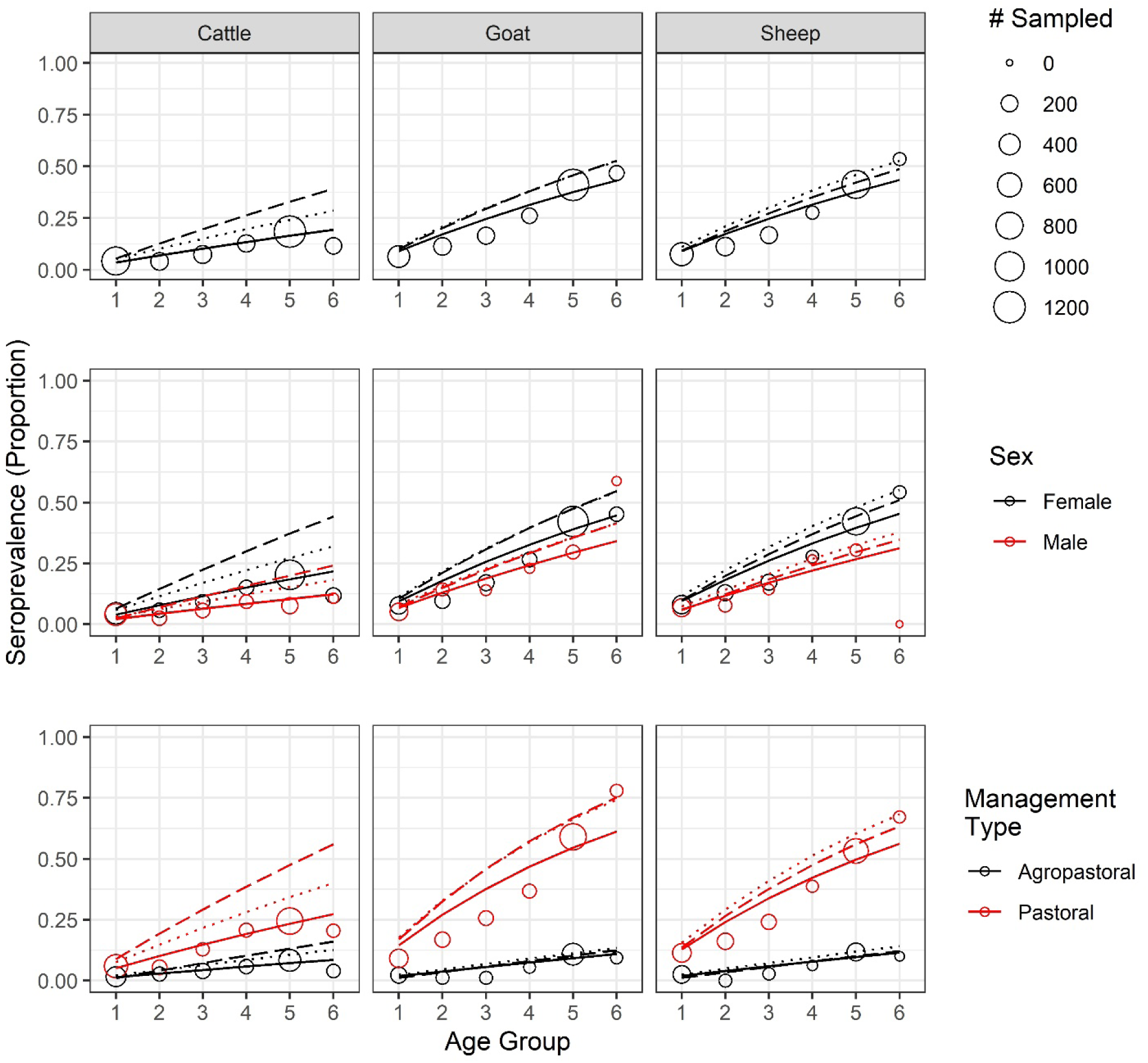
PPRV seroprevalence increases by age for sheep, goats, and cattle. Age-seroprevalence curves by species, sex, and management system. Solid lines indicate catalytic model fit to the apparent seroprevalence. True seroprevalence adjusted [50] for competitive ELISA antibody test sensitivity and specificity estimates of Couacy-Hymann et al 2007 [46] and Logan et al 2019 (unpublished) are plotted as dotted and dashed lines, respectively.

Age-seroprevalence curves, for both apparent and true seroprevalence (adjusted for cELISA test specificity and sensitivity, see Methods), are presented in Fig 2. Seroprevalence rose with age. In the oldest age cohort, observed seroprevalence for sheep, goats, and cattle reached 53.6%, 46.8%, 11.6%, respectively, while the highest adjusted, true seroprevalence reached 52.7%, 52.8%, and 39.2%, respectively. Both apparent and true sheep and goat seroprevalence was significantly different than cattle seroprevalence (p < 0.001). Notably, after adjustment the true cattle seroprevalence was 3.4 times the apparent cattle seroprevalence.

For most age groups (Fig 2, Table S3), females of any species had a higher apparent seroprevalence than males, except for male goats in the second and last age group; however, the only significant sex differences in apparent seroprevalence were among goats (p < 0.02) and cattle (p < 0.001) in age group 5. There were not enough male sheep (n=1) in the oldest age group to make comparisons, but the oldest male goats and cattle were significantly different from each other as were the oldest female sheep and goats when compared with cattle (p <0.001), but not with each other (p = 0.26). When adjusted, females had higher true seroprevalence across all ages than males.

Pastoral animals had higher apparent and true seroprevalence in each age group (Fig 2, Table S4), with the oldest animals reaching an apparent seroprevalence of 67.2%, 77.9%, and 20.5%, and highest true seroprevalence of 68.2%, 75.3%, and 56% in sheep, goats, and cattle, respectively. The oldest agropastoral animals reached an apparent seroprevalence of 10.0%, 9.4%, and 4.0%, and highest true seroprevalence of 14.0%, 13.5%, 16.1% in sheep, goats, and cattle, respectively. In pastoral systems, the oldest sheep and goats were significantly different from cattle (p << 0.001) but not from each other (p = 0.22). In agropastoral systems there was no significant difference between any pair of species (p > 0.28). Within each species, apparent seroprevalence was significantly different between management systems at each age group (p<0.05) with the exception of cattle in age group 2 (p = 0.44). After adjustment, all comparisons were significantly different. Strikingly, the oldest cattle seroprevalence estimate tripled in pastoral systems and quadrupled in agropastoral systems.

Nested models with different combinations of neighboring age intervals were compared to the maximal model of all five age groups (Fig S2) and the constant model of one FOI estimate across all ages (Tables S8-S10). The best fit models for each species are presented in Figs 4-6, with age-specific FOI estimates represented as a step function. For sheep, the best fit model had two age groups of 1-1.5 and 1.5-8 years, with the second age group having the highest FOI. For goats, the best fit model had three groups of 1-1.5, 1.5-5, and 5-8 years with the middle age group having the highest FOI and the first age group having the second highest. For cattle, the best fit model had four age groups of 1.5-2.5, 2.5-3.5, 3.5-4.5, and 4.5-10 years, with a clear FOI peak in the 3.5-4.5 age group, followed by the 2.5-3.5 age group. Profile confidence intervals (see Methods) around these best fitting models overlapped, indicating the FOI estimates for each age group were not significantly different. When stratified by sex (Fig 5), the best fit models are constant for males of all species as well as female sheep, but variable for female goats (3 age groups) and cattle (4 age groups). However, the confidence intervals for the sex- and age-specific FOI estimates overlapped within each of these models. When stratified by management system (Fig 6), the best fit models had varying numbers of age groups, all with overlapping FOI estimates, with the exception of pastoral cattle, which had a significantly different FOI estimate in the 2.3-3.5 year age group when compared to the other two age groups in the best fit model.

**Figure 4.**
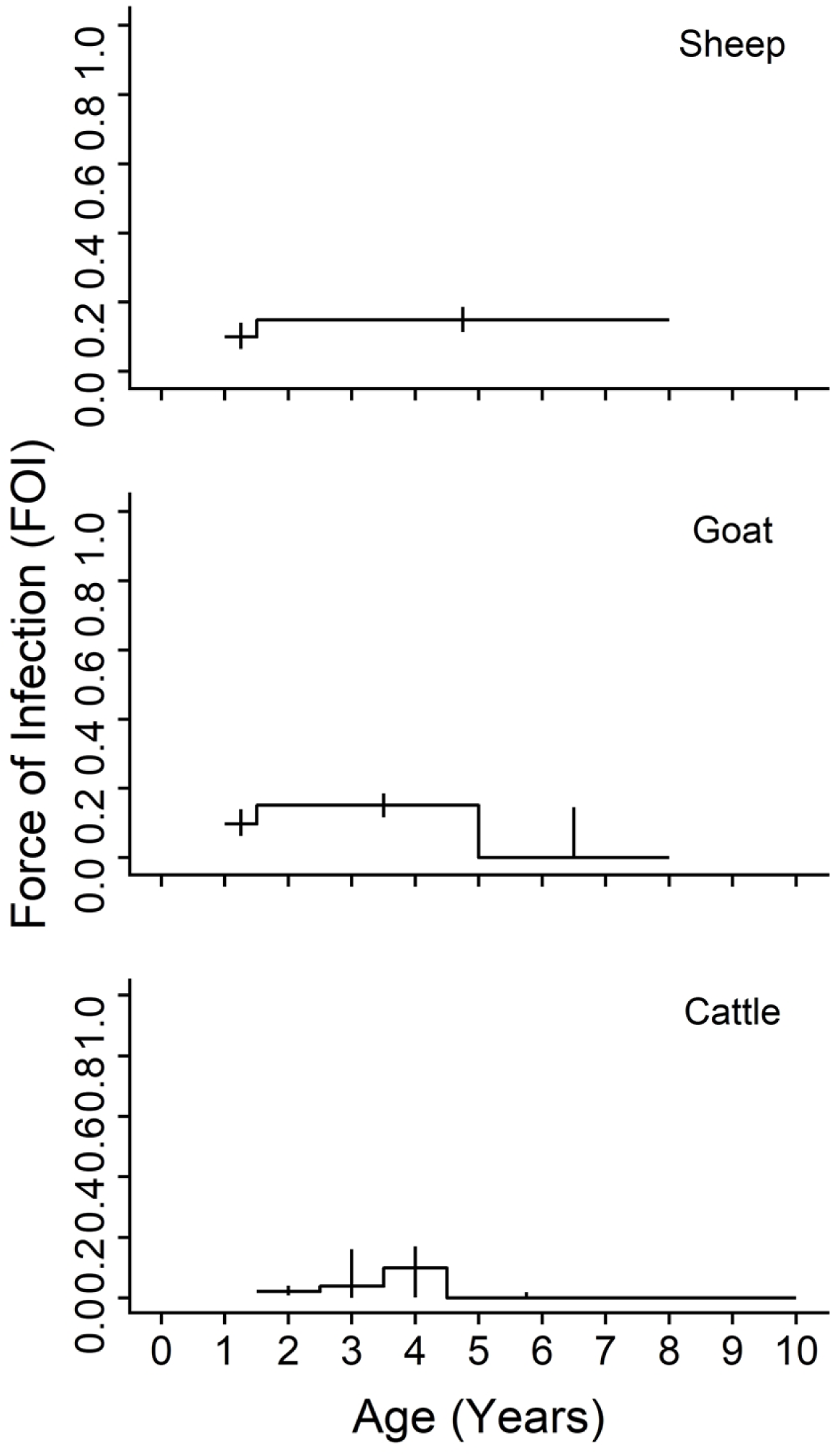
Age-varying force of infection models are a better fit than constant models for PPRV transmission by species. Age-specific force of infection estimates and profile confidence intervals from the best fit models for each species from a piece-wise catalytic model.

**Figure 5.**
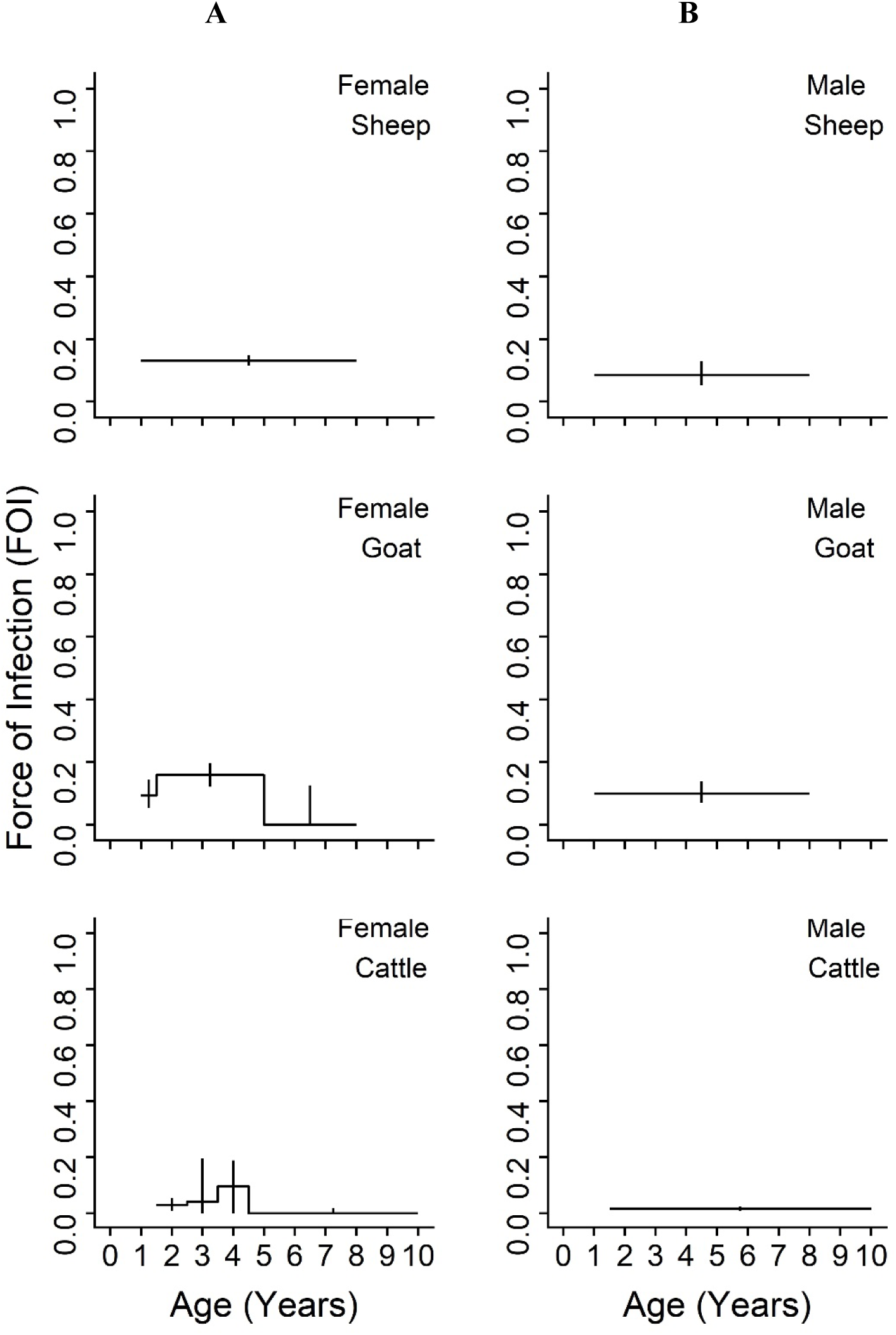
Age-varying force of infection models are a better fit for female goats and cattle than constant models for PPRV transmission by species and sex. Age-specific force of infection estimates and profile confidence intervals from the best fit models for each species from a piecewise catalytic model, stratified by sex. A) Females B) Males

**Figure 6.**
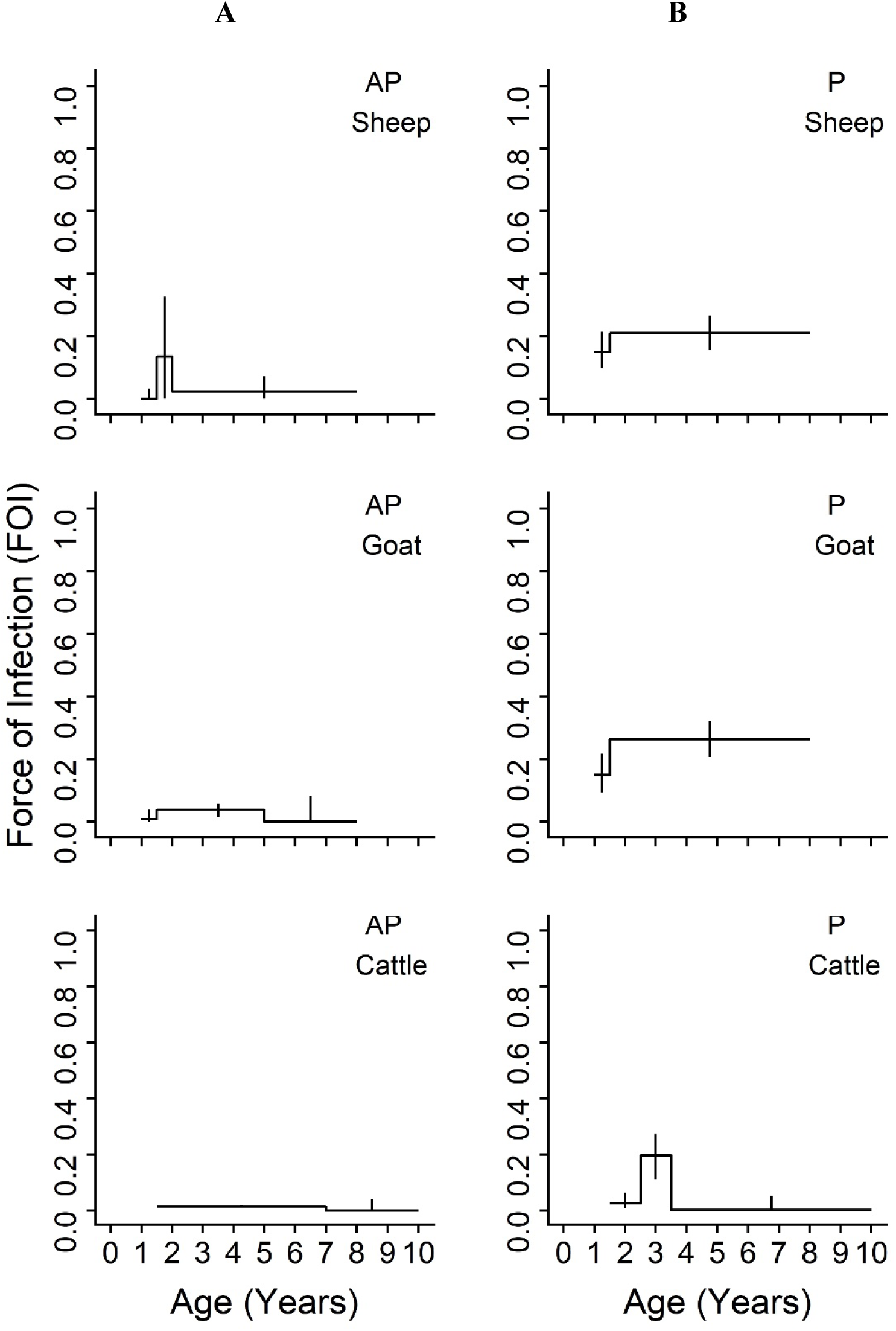
Age-varying force of infection models are a better fit than constant models for PPRV transmission by species and management system. Age-specific force of infection estimates and profile confidence intervals from the best fit models for each species from a piece-wise catalytic model, stratified by management system. A) Agropastoral (AP) B) Pastoral (P)

Lastly, logistic regression revealed that the impact of management system on PPRV seroconversion as measured by risk ratio was higher than all but the oldest age group (Fig S3), which had comparable impact when the confidence intervals were considered. The risk ratio for each species increased with age group as cumulative exposure increased over time.

## Discussion

Our study has demonstrated that i) PPRV seroprevalence increased across age for all species consistent with a pattern of endemic infection in which individuals’ cumulative exposure increases with age, ii) an age-varying model with a variable number of age groups by species provided a better fit to the data although age-specific FOI estimates were not significantly different from each other in most models, and iii) FOI estimates for all species were highest in younger age groups. PPRV seroprevalence was significantly higher in sheep and goats than cattle. Notably, the adjusted true cattle seroprevalence was triple the apparent seroprevalence (quadrupled in agropastoral, and tripled in pastoral systems). In the case of pastoral cattle, the adjustment resulted in true seroprevalence of 56%, which is greater than the highest reported cattle seroprevalence previously reported in the literature: 41.9% and 42% [20,21]. This result supports our previous finding [19] that cattle may play a more important role in PPRV transmission than previously realized, and also supports the importance of conducting more PPRV research in other species as called for by members of the PPR Global Research and Expertise Network (GREN) [4]. Across all ages, true PPRV seroprevalence was higher in females than in males, and higher in pastoral management systems than agropastoral systems. This was also true for the FOI for most age groups. The specific age patterns observed suggest a range of biological mechanisms for future study including the relationship between PPRV seroconversion and the age at first kidding, age at first market debut or mixing outside the herd, age at weaning, and age at waning of maternal immunity across production systems. Although the best fit models had age-varying FOI, the FOI estimates within most of these models were not significantly different due to the overlapping confidence interval estimates. This suggests that age may not play a strong role in PPRV transmission in the rural Tanzanian setting, with the exception of the significant signal observed in 2.5-3.5 year old pastoral cattle. Cattle and pastoral systems must be investigated in more detail to explain this finding. Taken together, the data presented here do not support targeted control by age group, instead they suggest that targeted control based on other risk factors, such as management system type, may be more effective [19].

These data suggest that all species experience higher FOI estimates when entering peak reproductive ages. Breeding in Tanzania tends to be uncontrolled, and the biological milestones of age at first breeding or kidding for sheep and goats has been estimated to range from 13.6-16.8 months [22,23] and for cattle 44.6–48 months [24,25], with average kidding intervals of 8-12 months for sheep and goats [22,26] and 12-26 months for cattle [24,25]. These estimates align well with FOI peaks seen in our maximal (Figs S1-S2) and best fit models (Figs 4-6). Outside of Tanzania, Dereje et al [27] reviewed several indigenous breeds in Ethiopia and across Africa, and found age at first kidding to range between 9-24 months with kidding interval between 7.9-12 months for goats depending on breed, location, and management system, with the earlier age at first kidding occurring more often in extensive, traditional management systems. Interestingly, the best fit models by sex are constant for males but variable for female goats and cattle suggesting potential subtle variation in PPRV transmission risk due to sex. This may possibly be due to within-host factors such as increased female susceptibility to infection due to immune changes during pregnancy (i.e. the periparturient response). However, it seems most likely that non-reproductive behavioral and management factors that lead to increased mixing at reproductive ages may drive the majority of age and sex PPRV transmission patterns observed. Increased mixing could increase transmission through the following mechanisms: i) interactions with animals at market where they are taken for sale, and sometimes return, at specific ages or maximum reproductive potential (2-4 years in Tanzanian settings, S. Cleaveland personal communication 2019), or ii) limited mixing of the youngest animals with older animals as part of husbandry practice (e.g. grazing or confinement overnight), or iii) husbandry mating system practices (e.g. increased temporary interaction with breeding male animals for servicing the herd), or iv) increased interaction with young animals who, after losing maternal immunity and prior to vaccination, become infected and pass PPRV to their mothers or other milking females they access prior to weaning between 3 and 6 months [28,29] (aligning with the higher seroprevalence and FOI estimates seen in females in this study).

Another important age-specific signal considered was maternal immunity. Maternal immunity to PPRV declines dramatically after birth, falling below the protective level by three [13,17], four [15], or five [14] months of age (note: there is not literature consensus on the protective level). Animals in this study were at least six months of age, so maternal immunity was not expected to play a role in the FOI patterns presented. Nevertheless, a six age group model (Fig S1, Tables S5-S7) and five age group model (Fig S2, Tables S8-S10) were explored, but neither were selected as the best fitting model among the nested models. FOI differences between these two models may be due to i) age misclassification among the youngest age group (i.e. they were younger than assessed) or ii) extended maternal immunity beyond the average weaning age (sheep and goats: 5 months [26], cattle: 7 months [24]) due to prolonged weaning or a differential length of maternal immunity provided by naturally infected mothers (expected in our sample) versus vaccinated mothers [13,14,35]. Future experiments are needed to clarify the role of immunity type and breed on maternal immunity duration as well as to examine PPRV transmissibility to young via milk [36] or reproductive tract secretions.

The finding that the apparent and true age-seroprevalence curves did not rise above 60% in the oldest age group when stratified by species or species and sex and is unexpected for a fully immunizing infection in an endemic setting. When stratified by management system, pastoral sheep and goats age-seroprevalence curves rose to the high proportion expected in endemic settings, whereas the curves among agropastoral animals did not, suggesting PPRV may not be endemic in agropastoral production systems (in agreement with PPRV modelling studies in Ethiopia [37]) and that management system differences explain the unexpected slow rise observed when stratifying only by species, or species and sex. Age-seroprevalence curves may also be affected by infection-associated mortality in sheep and goats, which would cause left truncation in our cross-sectional dataset and underestimation of the FOI (possibly more pronounced the smaller the FOI [38]). All animals sampled in this study were clinically healthy and no case fatality data were available to correct for infection-associated mortality among sheep and goats. Cattle are not known to suffer mortality from PPRV (although clinical disease and mortality have been seen Asian water buffalo [39], a non-primary host), so this effect is not expected to impact cattle age-seroprevalence curves, though more investigation is likely needed here. Currently, the available herd-level case fatality data in endemic settings is limited and variance is wide, which has been attributed to factors such as strain, species, and breed [18]. Where possible, future studies should strive to collect detailed case fatality data from recent or current epidemics in the study area to enable improved seroprevalence and FOI estimates in future cross-sectional studies.

Our study made several assumptions including: PPRV endemicity in Tanzania; age-stratified mixing of animals; a positive cELISA result indicates past PPRV exposure and current protection; a negative cELISA result indicates no past exposure and current susceptibility; that the BDSL cELISA kit was suitable for testing cattle samples; and that cross-reactivity with rinderpest or rinderpest-like viruses [40] was not expected in our samples given the sampling date and methods used to develop the cELISA kit. These assumptions (except mixing) are the same as those for the catalytic model with the assumption of constant age and have been discussed as reasonable first approximations to understanding PPRV transmission dynamics in previous work [19]. Additionally, while the analytical approach taken in this study can identify which age cohorts are responsible for the majority of transmission, it cannot distinguish whether PPRV infection is occurring mainly within a given age class or whether it originates from a different age class [41]. Misclassification of age in this study may have increased variation in FOI estimates in the last age group, by classifying more animals in the second to last age group (full mouth) than the last age group (full mouth plus wear) due to the difficulty of assessing age by variation in dental wear in full mouthed animals.

This study has demonstrated that the force of infection of PPRV varies significantly by age for pastoral cattle and that non-significant yet age-varying force of infection patterns are present among sheep, goats, and cattle. Management practices and biological milestones that occur near the identified force of infection peak ages should be targeted for further study. These data do not support targeted control by age group. Instead they suggest that targeted control based on management system may be more effective to achieve PPRV eradication.

## Methods

Blood samples were collected in 2016 from clinically healthy sheep, goats, and cattle in 20 northern Tanzanian villages in Arusha and Manyara Regions as part of the “Social, Economic, and Environmental Drivers of Zoonoses” (SEEDZ) study. The SEEDZ study data collection, data cleaning, and laboratory testing methods have been described in detail previously [19]. Relevant methods have been summarized here. Briefly, a total of 7,576 animals from 417 households and 404 household surveys were conducted using multistage random sampling. These households were from villages classified by livelihood type as either ‘pastoral’ (P) villages, which were those in which livestock rearing was considered the primary livelihood activity, or ‘agropastoral’ (AP), which were those villages in which a mix of crops and livestock were important livelihood activities. A maximum of 10 cattle, 10 sheep, and 10 goats per selected household were randomly selected for sampling as they moved through a crush. Among cattle, five animals between 6-18 months of age and five animals over 18 months of age were selected. Among sheep and goats, five animals between 6-12 months of age, and five animals over 12 months were selected. At least two of the animals were male in each of the groups, and all animals were old enough to be expected to lack maternally derived PPRV immunity [13].

Animal age was assessed by dentition, namely incisor tooth eruption and wear [42–44]. This age estimation method was used because exact recorded ages were not available. Among samples containing complete species, sex, age, and location data, a total of 7,538 serum samples were tested in duplicate using a commercial competitive ELISA kit (Pirbright Institute, Surrey, England) directed against the hemagglutinin protein of PPRV [45,46]. Samples were heat inactivated (56° C, 2 hours) prior to shipment to the University of Glasgow for testing. Forty-two samples (0.6% of total) were removed from analysis as they were from households that self-reported PPRV vaccination in the past 24 months. However, 427 samples (5.7% of total) were retained that lacked a household survey from which to discern self-reported PPRV vaccination as exclusion of these samples yielded qualitatively and quantitatively similar results. Our final analysis sample included 7,496 animals (2,080 sheep, 2,419 goats, 2,997 cattle).

To understand the role of age heterogeneities in PPRV transmission, we calculated the force of infection (FOI), the rate of infection of susceptible hosts, using the catalytic framework [47]. The catalytic model provides the framework for calculating the FOI from age-specific seroprevalence data from cross-sectional surveys [47–49]. Given that the rate at which immunity builds up with age depends on rates of circulation, this approach allows us both to elucidate the rate of circulation (FOI, λ) and to determine which age classes are most important for continued transmission. According to the catalytic model, age-specific seroprevalence, P(*a*), can be expressed as:

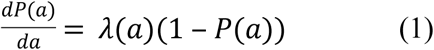

where (1-P(*a*)) is the proportion of susceptible hosts of age *a* and λ(*a*) is the age-specific FOI. Integrating and rearranging yields:

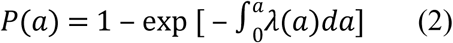

which describes how the cumulative FOI up to a given age will deplete a susceptible cohort, where P(*a*) represents the probability of having been infected before age *a*. Age-specific seroprevalence curves are direct empirical observations on this probability. This model assumes endemicity, age-stratified mixing, and that seronegatives are fully susceptible.

In this study, age-seroprevalence curves were used to describe the apparent and true (adjusted) seroprevalence patterns across ages. Adjustment was conducted according to the method of Rogan & Gladen 1978 [50], using recalculated sensitivity and specificity estimates from Couacy-Hymann et al [46], and new estimates from Logan et al (unpublished, personal communication 2019), who estimated specificity of > 99% for all three species, and sensitivity of 88%, 81%, and 48% in sheep, goats, and cattle, respectively, when cELISA was compared to the gold standard virus neutralization test. Power analysis indicated that a minimum sample size of 15 animals was needed in any age group to detect a 10% difference (medium effect size) between groups at power ≥ 80%. All stratifications except male sheep in age group 6 reached this criterion and a two proportion significance test was used to identify significant differences at a 5% level.

The piece-wise catalytic model (2) was used to calculate age-specific FOI by species, species and sex, and species and management system. The model assumes a fixed FOI within the specified age intervals (dentition-based age groups converted into age in years). The age intervals were defined by the upper age cutoff: 1, 1.5, 2, 3, 5, and 8 years for sheep and goats [42,43] and 1.5, 2.5, 3.5, 4.5, 7, and 10 years for cattle [44]. All individual animal ages were set to the midpoint of the age interval (e.g. an animal in dentition-based age group 1, with age interval 0 to 1 years, would have its age reset to 0.5 years).

The model was optimized in a three step process. Arbitrary initial parameter values (0.001) and a user defined function containing the model evaluated using exact integration were provided to the R software optim function. Parameter estimates were obtained using the quasi-Newton Broyden–Fletcher–Goldfarb–Shanno (BFGS) algorithm to minimize the negative log-likelihood. The first estimates were provided as initial values for a second optimization using the BGFS algorithm, and then the second estimates were used during a third optimization using the derivative free, stochastic, global optimization algorithm simulated annealing (SANN). Standard errors were computed using partial profile likelihood, following the method presented in Long et al [41]. Although all animals were six months or older and not expected to have maternal immunity [13], models with and without the first age interval (<1 year old, temporary teeth) were explored to demonstrate the impact on the FOI estimates. A model with five age intervals (first age interval removed) was preferred (presented in Fig S2, Tables S8-10) over the six age interval model (presented in Fig S1, Tables S5-S7), due to unrealistically low estimates of the second age interval. Additionally, nested models were used to understand the impact of different combinations of merged, neighboring age intervals on model fit. The best fitting model for each species was selected by Akaike Information Criterion [51] and are presented in the text. All analyses were conducted in R software version 3.5.3 [52].

## Ethics Statement

All adult participants in the SEEDZ study provided written informed consent. No data were collected from minors or children. In this study, questionnaire data were used solely to determine the PPRV vaccination status of household herds in the past 12 months. The protocols, questionnaire, and consent procedures were approved by the following ethical review committees: Kilimanjaro Christian Medical Centre (KCMC/832) and National Institute of Medical Research (NIMR/2028) in Tanzania; College of Medical, Veterinary and Life Sciences, University of Glasgow in the United Kingdom (200140152). Approval for animal work was provided by the Clinical Research Ethics Committee at the University of Glasgow School of Veterinary Medicine (39a/15), which authorizes research under The Veterinary Surgeons Act UK 1996 and oversees research regulated under the Animal (Scientific Procedures) Act 1986. Permission to publish this work was granted by Director of Veterinary Services, Tanzania.

## Acknowledgements

The authors thank all members of the SEEDZ field and laboratory team for their role in data acquisition and curation, including Tito Kibona, Kunda Mnzava, Euphrasia Mariki, Zanuni Kweka, Rigobert Tarimo, Fadhili Mshana, Mamus Toima, Tauta Maapi, Raymond Mollel, Matayo Lazaro, Ephrasia Hugo, Nelson Amana, Elizabeth Kasagama and Victor Moshi.

## Author Contributions

CMH, ONB conceptualized the study and methodology, designed and reviewed analyses. CMH conducted laboratory testing and formal analyses. WdG, BJW, SC, ESS provided additional data and metadata, laboratory resources, and advice. CMH wrote original manuscript, created visualizations including figures and tables. WdG, BJW, IMC, VK, PJH, JB, ESS, SC, ONB reviewed analyses and provided edits to manuscript.

## Data Availability

The anonymized dataset used in this study will be made available on the following repository: http://researchdata.gla.ac.uk/

## Supporting Information

**Supplemental Table 1.** Sample population characteristics, modified from Herzog et al 2019 [19]

**Supplemental Table 2.** Sample population distribution and apparent PPRV seroprevalence by dentition-based age group

**Supplemental Table 3.** Sample population distribution and apparent PPRV seroprevalence by dentition-based age group and sex

**Supplemental Table 4.** Sample population distribution and apparent PPRV seroprevalence by dentition-based age group and management system

**Supplemental Table 5.** Sheep nested models and AIC values for models combining five age groups

**Supplemental Table 6.** Goat nested models and AIC values for models combining five age groups

**Supplemental Table 7.** Cattle nested models and AIC values for models combining five age groups

**Supplemental Table 8.** Sheep nested models and AIC values for models combining six age groups

**Supplemental Table 9.** Goat nested models and AIC values for models combining six age groups

**Supplemental Table 10.** Cattle nested models and AIC values for models combining six age groups

**Supplemental Figure 1.** Age-specific force of infection estimates from the best fit piece-wise catalytic model with six age groups and age-seroprevalence curves by species. The model fit is plotted as a line, age group seroprevalence estimates as points, and the age-specific FOI estimates as a step function.

**Supplemental Figure 2.** Age-specific force of infection estimates from the best fit piece-wise catalytic model with five age groups and age-seroprevalence curves by species. The model fit is plotted as a line, age group seroprevalence estimates as points, and the age-specific FOI estimates as a step function.

**Supplemental Figure 3.** Logistic regression estimates of the impact of age, management, and sex on PPRV seroconversion. Reference group: male agropastoral cattle in age group 1. Management has a greater risk ratio and greater impact than all but the oldest age group(s).

**Supplemental Text.** Additional References that Tested the Significance of Age for PPRV Seroprevalence

## References

1. Global control and eradication of peste des petits ruminants. World Organization for Animal Health & Food and Agricultural Organization of the United Nations; 2015.

2. Zahur A Bin, Irshad H, Ullah A, Afzal M, Latif A, Ullah RW, et al. Peste des Petits Ruminants Vaccine (Nigerian Strain 75/1) Confers Protection for at Least 3 Years in Sheep and Goats. J Biosci Med. 2014;2:27–33.

3. Rossiter P. A Global PPR Network for Field Staff. Front Vet Sci [Internet]. 2019;6(August):1–4. Available from: https://www.frontiersin.org/article/10.3389/fvets.2019.00267/full

4. Baron MD, Diop B, Njeumi F, Willett BJ, Bailey D. Future research to underpin successful peste des petits ruminants virus (PPRV) eradication. J Gen Virol [Internet]. 2017;98(11):2635–44. Available from: http://www.microbiologyresearch.org/content/journal/jgv/10.1099/jgv.0.000944.v1

5. Woolhouse, MEJ., Dye, C., Etard, J-F., Smith, T., Charlwood, JD., Garnett, P., Hagan, P., Hii, JLK., Ndhlovu, PD., Quinnell, RJ., Watts, CH., Chandiwana, SK., Anderson R. Heterogeneities in the transmission of infectious agents: Implications for the design of control programs. Proc Natl Acad Sci. 1997;94(January):338–42.

6. Anderson RM, May RM. Vaccination and herd immunity to infectious diseases. Nature. 1985;318(6044):323–9.

7. Anderson RM, May RM. Directly Transmitted Infectious Diseases: Control by Vaccination. Science (80-). 1982;215(4536):1053–60.

8. Simon AK, Hollander GA, McMichael A. Evolution of the immune system in humans from infancy to old age. Proc R Soc B. 2015;282(20143085).

9. Hoover EA, Olsen RG, Hardy WD, Schaller JP, Mathes LE, Cockerell GL. Biologic and immunologic response of cats to experimental infection with feline leukemia virus. Bibl Haematol. 1976;No.43(2):180–3.

10. Fromont E, Artois M, Langlais M, Courchamp F, Pontier D. Modelling the feline leukemia virus (FeLV) in natural populations of cats (Felis catus). Theor Popul Biol. 1997;52(1):60–70.

11. Heisey DM, Joly DO, Messier F. The Fitting of General Force of Infection Models To Wildlife Disease Prevalence Data. Ecology. 2006;87(9):2356–65.

12. Elarbi AS, Kane Y, Metras R, Hammami P, Ciss M, Beye A, et al. PPR Control in a Sahelian Setting: What Vaccination Strategy for Mauritania? Front Vet Sci. 2019;6(242):1–18.

13. Bodjo C, Couacy-Hymann E, Koffi M, Danho T. Assessment of the duration of maternal antibodies specific to the homologous peste des petits ruminants vaccine Nigeria 75/1 in Djallonke lambs. Biokemistri. 2006;18(2):99–103.

14. Awa DN, Ngagnou A, Tefiang E, Yaya D, Njoya A. Post vaccination and colostral peste des petits ruminants antibody dynamics in research flocks of Kirdi goats and Foulbe sheep of north Cameroon. Prev Vet Med. 2002;55(4):265–71.

15. Balamurugan V, Sen A, Venkatesan G, Rajak KK, Bhanuprakash V, Singh RK. Study on passive immunity: Time of vaccination in kids born to goats vaccinated against peste des petits ruminants. Virol Sin. 2012;27(4):228–33.

16. Taylor WP. The Distribution and Epidemiology of Peste des Petits Ruminants. Prev Vet Med. 1984;2:157–66.

17. Obi T, Ojo M, Taylor W, Rowe L. Studies on the Epidemiology of Peste des petits ruminants in Southern Nigeria. Trop Vet. 1983;1:209–17.

18. Munir M, editor. Peste des Petits Ruminants Virus. Vol. I. Heidelberg: Springer; 2015.

19. Herzog CM, Glanville WA De, Willett BJ, Kibona TJ, Cattadori IM. Pastoral production is associated with increased peste des petits ruminants seroprevalence in northern Tanzania across sheep, goats and cattle. Epidemiol Infect. 2019;147(e242):1–9.

20. Khan HA, Siddique M, Abubakar M, Ashraf M. The detection of antibody against peste des petits ruminants virus in sheep, goats, cattle and buffaloes. Trop Anim Health Prod. 2008;40(7):521–7.

21. Ali WH, Osman NA, Asil RM, Mohamed BA, Abdelgadir SO, Mutwakil SM, et al. Serological investigations of peste des petits ruminants among cattle in the Sudan. Trop Anim Health Prod. 2018;51(3):655–9.

22. Chenyambuga S, Lekule F. Breed preference and breeding practices for goats in agropastoral communities of semi-arid and sub-humid areas in Tanzania. Livest Res Rural Dev. 2014;26(6).

23. Safari J, Mtenga L, Eik L, Sundstol F, Johnsen F. Analysis of three goat production systems and their contribution to food security in semiarid areas of Morogoro, Tanzania. Livest Res Rural Dev. 2008;20(5).

24. Kasonta J. Comparative growth and reproductive performance of Boran, Tanzania Shorthorn Zebu and Mpwapwa cattle breeds and various crosses. 1992.

25. Msanga Y, Mwakilembe P, Sendalo D. The indigenous cattle of the Southern Highlands of Tanzania: distinct phenotypic features, performance, and uses. Livest Res Rural Dev. 2012;24(7).

26. Chenyambuga S, Komwihangilo D, Jackson M. Production performance and desirable traits of Small East African goats in semi-arid areas of Central Tanzania. Livest Res Rural Dev. 2012;24(7).

27. Dereje T, Mengustu U, Getachew A, Yoseph M. A review of productive and reproductive characteristics of indigenous goats in Ethiopia. Livest Res Rural Dev. 2015;27(2).

28. Hyera E, Mlimbe ME, Sanka JD, Minja MG, Rugaimukamu AP, Latonga PM, et al. Onstation growth performance evaluation of Small East African and dual purpose goat breeds in Northern Tanzania. Livest Res Rural Dev. 2018;30(9).

29. Zergaw N, Dessie T, Kebede K. Growth performance of Woyto-Guji and Central Highland goat breeds under traditional management system in Ethiopia. Livest Res Rural Dev. 2016;28(1).

30. EFSA. Scientific Opinion on peste des petits ruminants. EFSA J. 2015;13(1):1–94.

31. OIE. Rinderpest [Internet]. OIE Technical Disease Card. 2013. Available from: http://www.oie.int/wahis/public.php?page=home]

32. Ladewig J, Price EO, Hart BL. Flehmen in male goats: Role in sexual behavior. Behav Neural Biol. 1980;30(3):312–22.

33. Banks E. Some Aspects of Sexual Behavior in Domestic Sheep, Ovis aries. Behaviour. 1964;23(3/4):249–79.

34. Chenoweth PJ. Sexual Behavior of the Bull: A Review. J Dairy Sci [Internet]. 1983;66(1):173–9. Available from: http://dx.doi.org/10.3168/jds.S0022-0302(83)81770-6

35. Markus TP, Adamu J, Kazeem HM, Olaolu OS, Woma TY. Assessment of Peste des petits ruminants antibodies in vaccinated pregnant Kano brown does from Nigeria and subsequent maternal immunity in their kids. Small Rumin Res. 2019;174:53–6.

36. Clarke BDBD, Islam MRMR, Yusuf MAMA, Mahapatra M, Parida S. Molecular detection, isolation and characterization of Peste-des-petits ruminants virus from goat milk from outbreaks in Bangladesh and its implication for eradication strategy. Transbound Emerg Dis. 2018;(April):1597–604.

37. Fournié G, Waret-szkuta A, Camacho A, Yigezu LM, Pfeiffer DU. A dynamic model of transmission and elimination of peste des petits ruminants in Ethiopia. Proc Natl Acad Sci. 2018;115(33):8454–9.

38. Caley P, Hone J. Estimating the force of infection; Mycobacterium bovis infection in feral ferrets Mustela furo in New Zealand. J Anim Ecol. 2002;71(1):44–54.

39. Govindarajan R, Koteeswaran A, Venugopalan AT, Shyam G, Shaquna S, Shaila MS, et al. Isolation of pestes des petits ruminants virus from an outbreak in Indian buffalo (Bubalus bubalis). Vet Rec. 1997;141(22):573–4.

40. Logan N, Dundon WG, Diallo A, Baron MD, Nyarobi MJ, Cleaveland S, et al. Enhanced immunosurveillance for animal morbilliviruses using vesicular stomatitis virus (VSV) pseudotypes. Vaccine [Internet]. 2016;34(47):5736–43. Available from: http://dx.doi.org/10.1016/j.vaccine.2016.10.010

41. Long GH, Sinha D, Read AF, Pritt S, Kline B, Harvill ET, et al. Identifying the age cohort responsible for transmission in a natural outbreak of Bordetella bronchiseptica. PLoS Pathog. 2010;6(12):1–10.

42. Mitchell T. NSW Agriculture AGFACTS: to tell the age of goats [Internet]. 2003. Available from: https://www.dpi.nsw.gov.au/animals-and-livestock/goats/mgt/age

43. Casburn G. NSW Agriculture PrimeFact: How to tell the age of sheep [Internet]. Agriculture. 2016. Available from: https://www.dpi.nsw.gov.au/animals-and-livestock/sheep/management/general-information/age

44. Turton JA. How to estimate the age of cattle [Internet]. 1999. Available from: http://www.nda.agric.za/docs/Infopaks/cattleEstimateAge.pdf

45. Anderson J, McKay JA, Butcher RN. The use of monoclonal antibodies in competitive ELISA for the detection of antibodies to rinderpest and peste des petits ruminants viruses. IAEA-TECDOC-623 The sero-monitoring of rinderpest throughout Africa: phase I. 1991.

46. Couacy-Hymann E, Bodjo SC, Tounkara K, Koffi YM, Ohui a H, Danho T, et al. Comparison of two competitive ELISAs for the detection of specific peste-des-petits-ruminant antibodies in sheep and cattle populations. African J Biotechnol. 2007;6(6):732–6.

47. Hens N, Shkedy Z, Aerts M, Faes C, Van Damme P, Beutels P. Modeling infectious disease parameters based on serological and social contact data: A modern statistical perspective. New York: Springer; 2012.

48. Hens AN, Aerts M, Faes C, Shkedy Z, Lejeune O, Damme PVAN, et al. Seventy-five years of estimating the force of infection from current status data. Epidemiol Infect. 2010;138:802–12.

49. Muench H. Derivation of Rates from Summation Data by the Catalytic Curve. J Am Stat Assoc. 1934;29(185):25–38.

50. Rogan WJ, Gladen B. Estimating Prevalence from the Results of a Screening Test. Am J Epidemiol. 1978;107(1):71–6.

51. Akaike H. A New Look at the Statistical Model Identification. IEEE Trans Automat Contr. 1974;19(6):716–23.

52. R Core Team (2018). R: A language and environment for statistical computing. R Foundation for Statistical Computing, Vienna, Austria. URL https://www.R-project.org/.

